# Dynamic dysregulation of Tenascin-X/Tenascin-C balance controlled by Transforming Growth Factor-β leads to tumor cell proliferation during pancreatic carcinogenesis

**DOI:** 10.1101/2025.02.07.635757

**Authors:** Sophie Liot, Céline Schmitter, Perrine Mercier-Gouy, Naïma El Kholti, Jonathan Balas, Laura Prigent, Alexandre Aubert, Claire Lethias, Jessica L. Chitty, Manuel Koch, Tanguy Fenouil, Thomas R. Cox, Ana Hennino, Valérie Hervieu, Sylvie Ricard-Blum, Ulrich Valcourt, Elise Lambert

## Abstract

Pancreatic ductal adenocarcinoma (PDAC) is one of the deadliest cancers due to late diagnosis and poor therapeutic efficiency. Throughout PDAC progression, a dense extracellular matrix (ECM) is deposited around neoplastic cells and accompanies tumor development and aggressiveness. This significant stroma, both in terms of quantity and through its impact on tumor cells, is now considered as a target for innovative therapies to improve patient survival. Among ECM proteins, Tenascins (TNs) are a family of four glycoproteins (TNC, TNW, TNR and TNX) sharing a common modular structure, but exhibiting different expression patterns and functions depending on the physiological and physio-pathological contexts. In PDAC, TNC is up-regulated and is considered as a pro-tumorigenic actor, whereas TNX’s role remains to be elucidated. Herein, we demonstrated that unlike TNC, TNX is drastically decreased in PDAC, and that this loss is correlated with reduced patient survival. Dysregulation of the TNX/TNC balance is attributable to Transforming Growth Factor-beta (TGF-β) upregulation during pancreatic carcinogenesis. Interestingly, we found that TNX is first highly deposited in low grade lesions before being clearly decreased in later stages suggesting an elaborate stromal remodelling during PDAC development. Finally, low TNX and high TNC deposition around precursor lesions are correlated with increased preneoplastic cell proliferation. Altogether, our results (i) demonstrate the importance of Tenascin ratios during pancreatic carcinogenesis, (ii) suggest an anti-proliferative role for TNX unlike its pro-tumoral counterpart, TNC and (iii) underscore TNX and TNC as valuable targets for the development of new drugs for pancreatic cancer and/or solid tumor treatment.

**Statement of significance:** The tumor microenvironment plays a prominent but contradictory role in pancreatic cancer. Our study highlights the role of a TGF-β-controlled TNX/TNC balance and designates TNX as an inhibitor of tumor cell proliferation.

## Introduction

Pancreatic Ductal Adenocarcinoma (PDAC) represents about 90% of pancreatic cancers, and is one of the deadliest cancers worldwide, with a 5-year relative survival rate below 10% (1). This poor prognosis is a consequence of difficulty in diagnosis. Indeed, PDAC develops without specific symptoms, and patients are thus usually at an advanced stage of the disease at diagnosis, with locally advanced or metastatic tumors (2). As a consequence, only 20% of tumors are resectable and conventional chemotherapy is poorly effective, extending patient lifespan by only few months (3). Thus, pancreatic cancers were ranked as the sixth cause of cancer mortality worldwide in 2022 (https://gco.iarc.fr/, (4)), and are predicted to become the second leading cause of cancer-related deaths in industrialized countries by 2030, if no significant therapeutic progress is made.

PDAC classically develops from precursor lesions, the main ones being Pancreatic Intraepithelial Neoplasia (PanINs). These pre-neoplastic lesions progress according to a linear model, and are classified into different grades characterized by the evolution of epithelial cell morphology from low grade PanINs to high grade PanINs, also referred as *in situ* carcinoma (5). Invasive PDAC then arises after basement membrane degradation and invasion of the underlying tissue by tumor cells. PDAC development is accompanied by the accumulation of genetic alterations in (pre)-neoplastic cells, from *KRAS* oncogene activation, to tumor suppressor loss such as *CDKN2A*, *TP53* and *SMAD4*, and finally culminates with around 60 mutations documented in PDAC tumors (6).

PDAC is also characterized by an intense desmoplastic reaction around lesions, *i.e.* an excessive extracellular matrix (ECM) deposition due to the recruitment of activated fibroblasts originating from (i) resident fibroblasts, (ii) Pancreatic Stellate Cells (PSCs), or (iii) Mesenchymal Stem Cells (MSC) (7). Thus, the connective tissue surrounding pancreatic cells increases from 5% in normal pancreas, to about 60% of the tumor mass (up to 90% in some cases) (8). This dense stroma, which is also associated with hypovascularization, can act as a physical barrier to conventional chemotherapies, and has been correlated with poor prognosis (9). Initially ignored, the tumor microenvironment (TME) is now the subject of intense research, as it influences tumor progression and thus represents a new therapeutic target. However, therapeutic strategies based on stromal cell suppression have failed (10,11), revealing that the tumor stroma also contains tumor-restraining components and highlighting the high complexity of this particular compartment (12).

The ECM is a meshwork of proteins/glycoproteins, mostly composed of fibers (elastic and collagens), fibronectins, laminins and proteoglycans (13). While historically described as an inert support for cells with only an architectural role, the ECM is now known to influence cell behavior either (i) directly by interacting with cells, or (ii) indirectly by sequestrating cytokines and growth factors, that can be released in response to stimuli (13). This signaling role has been evidenced not only for constitutive ECM in healthy connective tissue, but also for the tumor-surrounding ECM. Indeed, many matrix proteins can modify cancer cell behavior, and thus be involved in cancer progression (14), as we recently reviewed for the particular case of PDAC (12). Matricellular proteins are a group of matrix proteins regulating cell behavior (15). Among them, Tenascins (TNs) are a group of 4 glycoproteins (TNC, TNR, TNW, TNX) sharing a common modular structure with an oligomerization domain involved in subunit trimerization through coil-coil interactions, repeats of EGF-like and Fibronectin III (FN-III) domains, and a C-terminal Fibrinogen-like globular domain (FBG) (16). However, TNs display very different spatio-temporal expression patterns in physiological and physio-pathological contexts, and their expression and roles in tumor progression have been unequally studied.

Tenascin-C (TNC) (long monomeric isoform=241kDa), which forms hexamers thanks to a disulfide bond between two trimers, is the most extensively studied. It is usually expressed during development, at the embryonic epithelial-mesenchymal interface, around motile cells, during branching morphogenesis, and in developing smooth muscle, bone and cartilage. In adults, TNC expression is restrained to some stem cell niches, tendons, myotendinous junctions and the nervous system (17). However, TNC can undergo *de novo* synthesis and deposition in several pathological contexts, such as carcinogenesis and TNC expression is correlated with poor prognosis in many cancers, including several digestive tract tumors, lung, prostate and breast cancers and melanoma (18). Tenascin-W (TNW, monomer=144kDa) expression has been far less studied, but is detected during embryogenesis, notably at sites of osteogenesis, and is usually restricted to the periosteum, kidney and some muscles in adults (19). Like TNC, its expression is increased in numerous solid tumors, including breast, colorectal, prostate, lung, ovarian and brain cancers and melanoma (20). Tenascin-R (TNR, monomer=150kDa) presents an even more restricted expression pattern, being almost exclusively expressed in the nervous system, during development and in adults (20). The status of TNR during tumor progression is more controversial, as TNR has been found overexpressed in some but not all brain tumors (20). Finally, Tenascin-X (TNX, long monomeric isoform=458kDa), the largest member of the TN family, encoded by *TNXB* gene, presents a very broad expression pattern. Indeed, it is expressed from late developmental stages, suggesting a role during organogenesis, and persists in numerous organs in adults (21). Within these tissues, TNX plays a dual role, with (i) a structural function as an organizer of the collagenous network, by regulating the spacing between collagen fibers through its quaternary structure, and (ii) a matricellular role, due to its modulation of cell adhesion, and the regulation of growth factor availability, notably that of TGF-β, through its fibrinogen-like (FBG) domain (22). Compared to TNC and TNW, the status and the role of TNX during tumor progression remains controversial (23–28). In a previous study, we analyzed the expression of TNX in many cancers, including the 6 most common cancers worldwide, and demonstrated that (i) TNX expression is downregulated in almost all studied cancers - except in brain tumors, in which TNX levels are increased - and (ii) low level of *TNXB* expression in lung and breast tumors is associated with reduced patient survival (29). In keeping with these findings, another study including 15 tumor types, demonstrated that *TNXB* is one of the most significantly downregulated genes in cancer samples compared to healthy tissue (30). In addition, it has been demonstrated that melanoma cells injected in the foot pad of *TNXB* knockout mice led to an increase in primary tumor size and lung metastases compared to wildtype mice (31). Altogether, these studies suggest TNX could have a protective role against tumor development and progression.

Herein, we aimed at determining the profile of TNX deposition in PDAC, and elucidating its role in tumor progression. To strengthen our study, TNX results were compared to that of TNC, which is up-regulated during pancreatic cancer development.

## Results

### TNX expression is decreased in human PDAC samples at both mRNA and protein levels

We determined *TNXB* gene expression in human PDAC samples using microarray datasets from the Gene Expression Omnibus (GEO) repository (https://www.ncbi.nlm.nih.gov/geo/) comprising total resected PDAC and normal tissue. We searched for datasets with more than 20 patients and paired controls (matched normal tissue adjacent to the tumor, NAT) to avoid inter-patient variability (Supplementary Table S1). Two cohorts meeting these criteria were selected, GSE62452 and GSE15471 (60 and 36 patients respectively), and compared with GEO2R (https://www.ncbi.nlm.nih.gov/geo/geo2r/) to identify differentially expressed genes (DEGs) between tumor and NAT samples. In these datasets, *TNXB* was found to be downregulated in tumor compared to NAT (Supplementary Tables S1, S3 and S4). We then extracted and plotted normalized expression values for the best probe corresponding to *TNXB* gene and compared both groups. *TNXB* mRNA level was significantly decreased (*p*<0.001) in PDAC samples compared to NAT samples in both cohorts (Figure 1A). Additionally, we determined the fold change between tumor and NAT tissue for each patient, and found that more than 75% of patients had a decreased *TNXB* expression in their tumors (Figure 1B).

**Figure 1:**
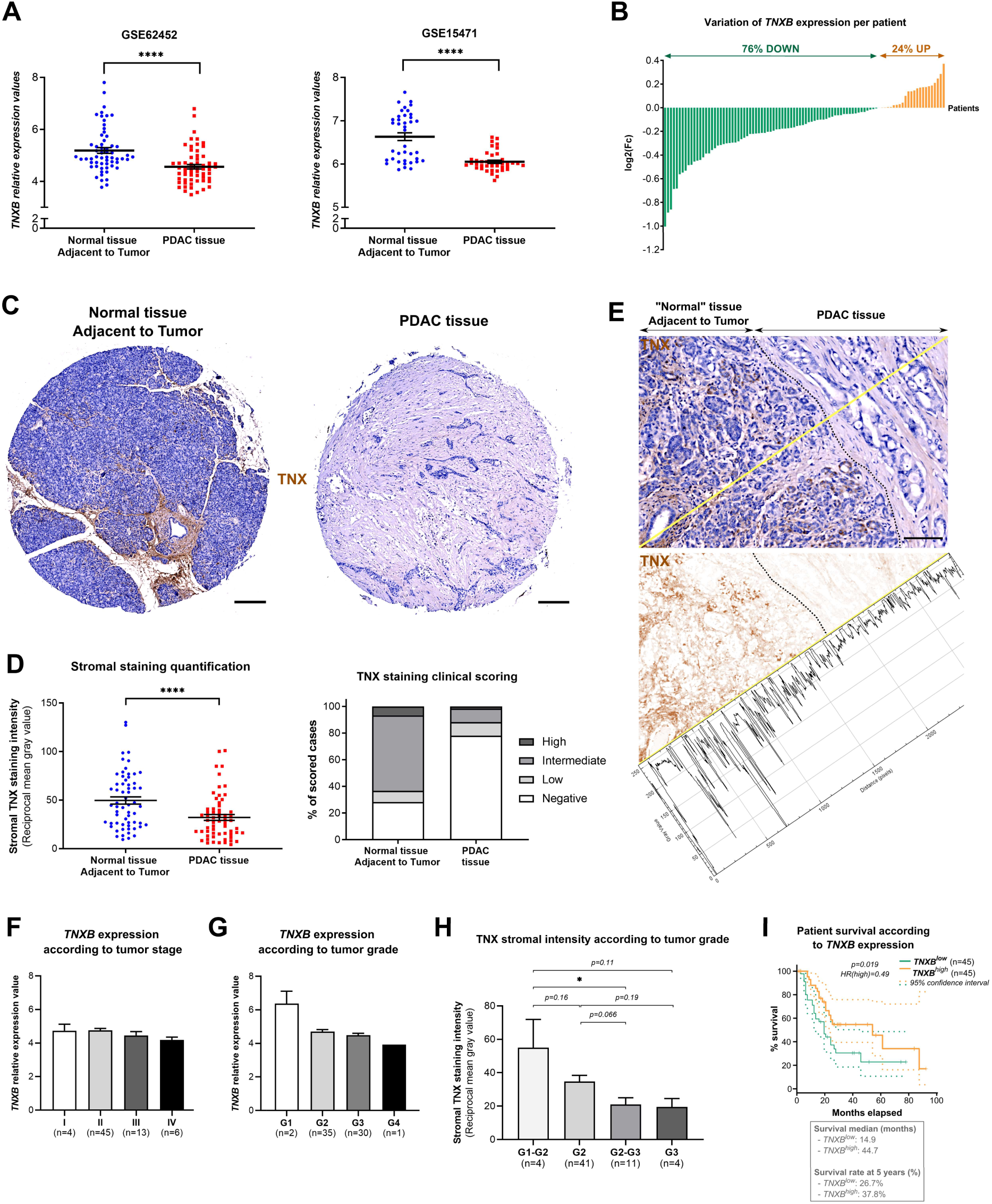
Tenascin-X loss in human PDAC samples predicts poorer prognosis. **A**- Normalized *TNXB* expression values for the two selected GEO datasets (individual values with mean ± SEM). **B**- Log2(Fold change) of *TNXB* expression values between PDAC and normal tissue adjacent to the tumor (NAT) for patients from both cohorts. Green and orange bars indicate negative and positive Log2(Fc), respectively. **C**- Representative images of NAT and PDAC cores immunolabelled with anti-TNX antibodies from analyzed TMA. Scale bar= 200 μm. **D**- Left: Quantitative analysis of TNX labelling in the stromal compartment of PDAC and NAT cores of TMA. Right: Clinical scoring of TNX staining in stromal part of PDAC and NAT cores of TMA by an anatomical pathologist. **E**- Top: Picture of a “PDAC” tissue core focused on both tumoral and non-tumoral zones following TNX immunolabelling and Gill’s hematoxylin counterstaining. Scale bar= 100 μm. Bottom: Same image with only DAB (TNX) staining and TNX quantification (histogram) on the yellow line. **F**- *TNXB* gene expression values in the samples of patients from the GSE62452 cohort according to AJCC stage. **G-** *TNXB* gene expression values in the samples of patients from the GSE62452 cohort according to WHO grade. **H-** Stromal TNX intensity in the cores of patients on the analyzed TMA according to the tumor grade (WHO). **F-H-** Number of patients in each group is indicated (n=). **I**- Cumulative survival proportion depending on high (orange) or low (green) *TNXB* transcript level in PDAC samples from patients using the GEPIA software. Survival median and survival rate at 5 years for each group of patients are indicated in gray.

As shown in Supplementary Table S1, only the two selected datasets showed significant differences in *TNXB* expression. However, 7 out of 8 datasets (87.5%) showed a decrease in *TNXB* expression even though it did not reach significance. This is likely due to the low number of samples per group (Supplementary Table S1). Indeed, when we performed a post-hoc power analysis using an Experimental Design Assistant tools (https://eda.nc3rs.org.uk/experimental-design-group#PowerCalc), we found that the minimal sample size to obtain significant difference should be 41 or 20 for GSE62452 or GSE15471 datasets, respectively. Additionally, although NAT samples appear to be histologically normal, their close vicinity to the tumor tissue might influence their expression profile, thus leading to misinterpretation. Therefore, we also analyzed other datasets from the GEO repository, with normal pancreas as controls, which allowed us to confirm a significant *TNXB* downregulation in tumor samples compared to normal pancreatic tissues in 57% of the datasets (Supplementary Table S2). It should be noted that the other datasets only led to statistically non-significant results which was probably due again to the low number of samples in normal or tumor categories.

At the protein level, TNX was immunodetected on a Tissue MicroArray (TMA) slide containing 80 human PDAC samples, 62 being associated with NAT samples (Figure 1C, supplementary Figure S1). We measured TNX level in the stroma of normal pancreas and tumor and observed a significant decrease of TNX staining in PDAC tissues, compared to adjacent non-tumor cores (Figure 1D, left). TMA slide was also analyzed by a pathologist specialized in pancreatic cancer, who scored each core into 4 categories: negative, low, intermediate or high TNX level. Almost 80% of PDAC cores were scored as negative, compared to 30% for NAT cores, for which 50% presented an intermediate level of TNX in their connective tissue (Figure 1D, right). Interestingly, in some cores initially classified as PDAC samples, TNX labelling was negative in the tumor compartment, while it was positive in the normal tissue adjacent to the tumor (Figure 1E). This was confirmed by plotting TNX staining intensity measured along a line passing through PDAC and adjacent tissues, and the intensity was indeed higher in NAT than in tumor tissue (Figure 1E). It should be noted that in the “healthy zone”, intense TNX staining was observed in cells clustered around islets that do not resemble classical pancreatic connective tissue, again suggesting that it has been altered due to the proximity to tumor cells. Altogether, our results clearly demonstrated that TNX level is drastically decreased in the stroma of human PDAC.

### Low *TNXB* expression is associated with shorter patient survival

To determine if the loss of TNX expression was correlated with disease progression, and if this could be deleterious for patients, we plotted *TNXB* expression values as a function of the American Joint Commission on Cancer (AJCC) stages (*i.e.* Tumor/Node/Metastasis (TNM) classification) or to World Health Organization (WHO) grades for each patient in the GSE62452 cohort. We did not observe a significant difference in *TNXB* expression level as a function of AJCC stages, which reflects the size and invasiveness of the tumor (Figure 1F). However, we clearly observed a decrease in *TNXB* expression depending on WHO grade progression, which reflects the differentiation of the tumor compared to normal tissue (Figure 1G). This result was further confirmed by analyzing stromal TNX protein levels depending on tumor grades in the TMA (Figure 1H). Again, due to the low number of samples in G1 and G3 or G4 tumors, both results regarding tumor grade only tended to be statistically different.

We next analyzed PDAC patient survival depending on *TNXB* mRNA level using Gene Expression Profiling Interactive Analysis (GEPIA) software (http://gepia2.cancer-pku.cn/). The Cancer Genome Atlas (TCGA)/Genotype-Tissue Expression (GTEx) data were divided into two groups: *TNXB*^high^ and *TNXB*^low^ patients corresponding to those with the highest and the lowest *TNXB* expression in their tumor (first and last quartiles), respectively. The Kaplan–Meier curve showed that low expression of *TNXB* (*TNXB*^low^) in tumors was significantly correlated with a poor prognosis for PDAC patients, compared to high expression of *TNXB* (*TNXB*^high^) (Figure 1I). The median survival was 44.7 months for *TNXB*^high^ patients *versus* 14.9 months for *TNXB*^low^ patients (Figure 1I). We thus demonstrated here that *TNXB* mRNA and TNX protein levels are markedly decreased in human PDAC, and that low *TNXB* expression is associated with poor prognosis for patients.

### TNC is dramatically increased in human PDAC samples, but is not a prognostic factor

TNC is upregulated in the stroma of most solid tumors and plays significant roles in proliferation, invasion, angiogenesis and metastasis (32). The analysis of *TNC* expression in the two selected cohorts showed that *TNC* mRNA levels were significantly higher in PDAC samples compared to “normal” tissues (Figure 2A) and was up-regulated in 70% of PDAC samples of patients compared to their NAT (Figure 2B). The same conclusions were also drawn at the protein level by immunodetecting TNC on the PDAC TMA slide. Indeed, TNC was deposited in higher amounts in the tumor area compared to NAT (Figure 2C-E, Supplementary Figure S1). It should be noted that in the zone where both “normal” and tumoral tissues were observed, we clearly detected TNC expression in the tissue adjacent to the tumor even though this area was classified as histologically “normal” by an anatomical pathologist suggesting that TNC production and deposition is an early event in pancreatic carcinogenesis occurring before the morphological modification of the pancreatic tissue (Figure 2E, Supplementary Figure S1). Also, TNX and TNC deposition on serial sections were consistently of a mutually exclusive pattern (Supplementary Figure S1). Despite the significant upregulation of TNC at both transcriptional and translational levels, it was not correlated with the stage/grade of the tumor or with patient survival (Figure 2F-I).

**Figure 2:**
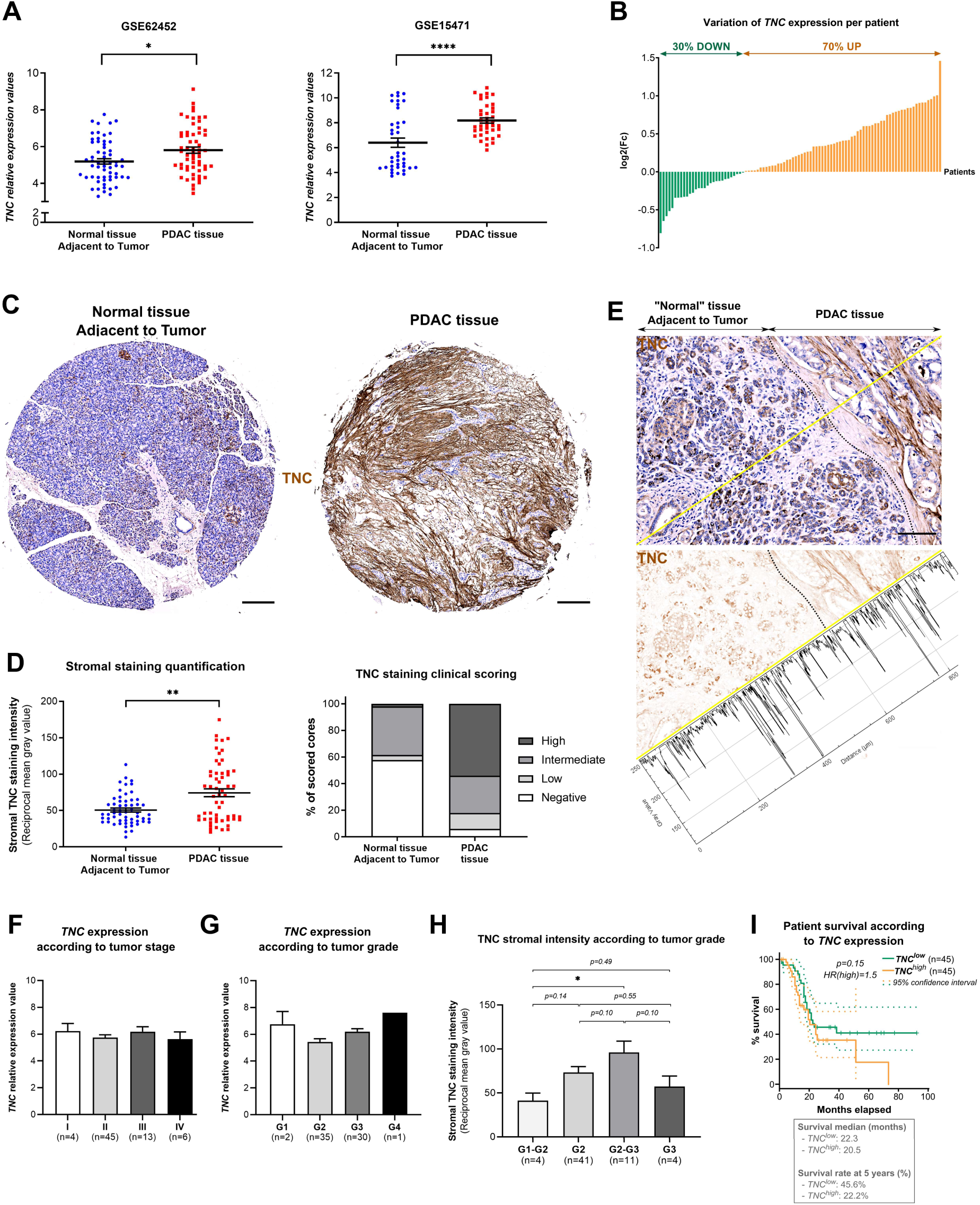
Tenascin-C is increased in human PDAC samples, but cannot be considered as a prognostic factor. **A**- Normalized *TNC* expression values for the two selected GEO datasets (individual values with mean± SEM). **B**- Log2(Fold change) of *TNC* expression values between PDAC and normal tissue adjacent to the tumor (NAT) for patients from both cohorts. Green and orange bars indicate negative and positive Log2(Fc), respectively. **C**- Representative images of NAT and PDAC cores immunolabelled with anti-TNC antibodies from analyzed TMA. Scale bar= 200 μm. **D**- Left: Quantitative analysis of TNC labelling in the stromal compartment of PDAC and NAT cores of TMA. Right: Clinical scoring of TNC staining in the stroma of PDAC and NAT cores by an anatomical pathologist. **E**- Top: Image of a “PDAC” tissue core focused on both tumoral and non-tumoral zones following TNC immunolabelling and Gill’s hematoxylin counterstaining. Scale bar= 100 μm. Bottom: Same image with only TNC immunostaining and TNC quantification (histogram) on the yellow line. **F**- *TNC* gene expression values in the samples of patients from the GSE62452 cohort according to AJCC stage. **G-** *TNC* gene expression values in the samples of patients from the GSE62452 cohort according to WHO grade. **H-** Stromal TNC intensity in the cores of patients on the analyzed TMA according to the tumor grade (WHO). **F-H-** The number of patients in each group is indicated (n=). **I**- Cumulative survival proportion depending on high (orange) or low (green) *TNC* transcript level in PDAC samples from patients using the GEPIA software. Survival median and survival rate at 5 years for each group of patients are indicated in gray.

### Opposite dysregulation of TNX and TNC is also observed in murine pancreatic tumors

To understand the roles of altered genes and to recapitulate the histological and molecular hallmarks observed during the different steps of pancreatic carcinogenesis from precursor lesions to PDAC, various genetically engineered mouse models (GEMMs) of PDAC have been developed over the past 20 years (33). We first analyzed *Tnxb* and *Tnc* expression in the aggressive [*Pdx1-Cre; LSL-Kras^G12D^; Ink4a/Arf^lox/lox^*] (KIC) mouse model using the GSE61412 dataset from the GEO repository. As expected from our results obtained using human samples, *Tnxb* transcripts were significantly downregulated in the PDAC tissue of KIC mice compared to the pancreas of normal KI mice whereas *Tnc* mRNA expression was upregulated (Figure 3A). The immunodetection of TNX and TNC on pancreatic sections demonstrated a loss of TNX in PDAC of 7-8-week-old KIC mice compared to age-matched WT mice, while TNC was abundantly deposited in the tumor tissue (Figure 3B and C). These results, in the KIC mouse model, correlate with the patient sample cohorts and again highlight the opposite dysregulation of TNX and TNC in PDAC samples.

**Figure 3:**
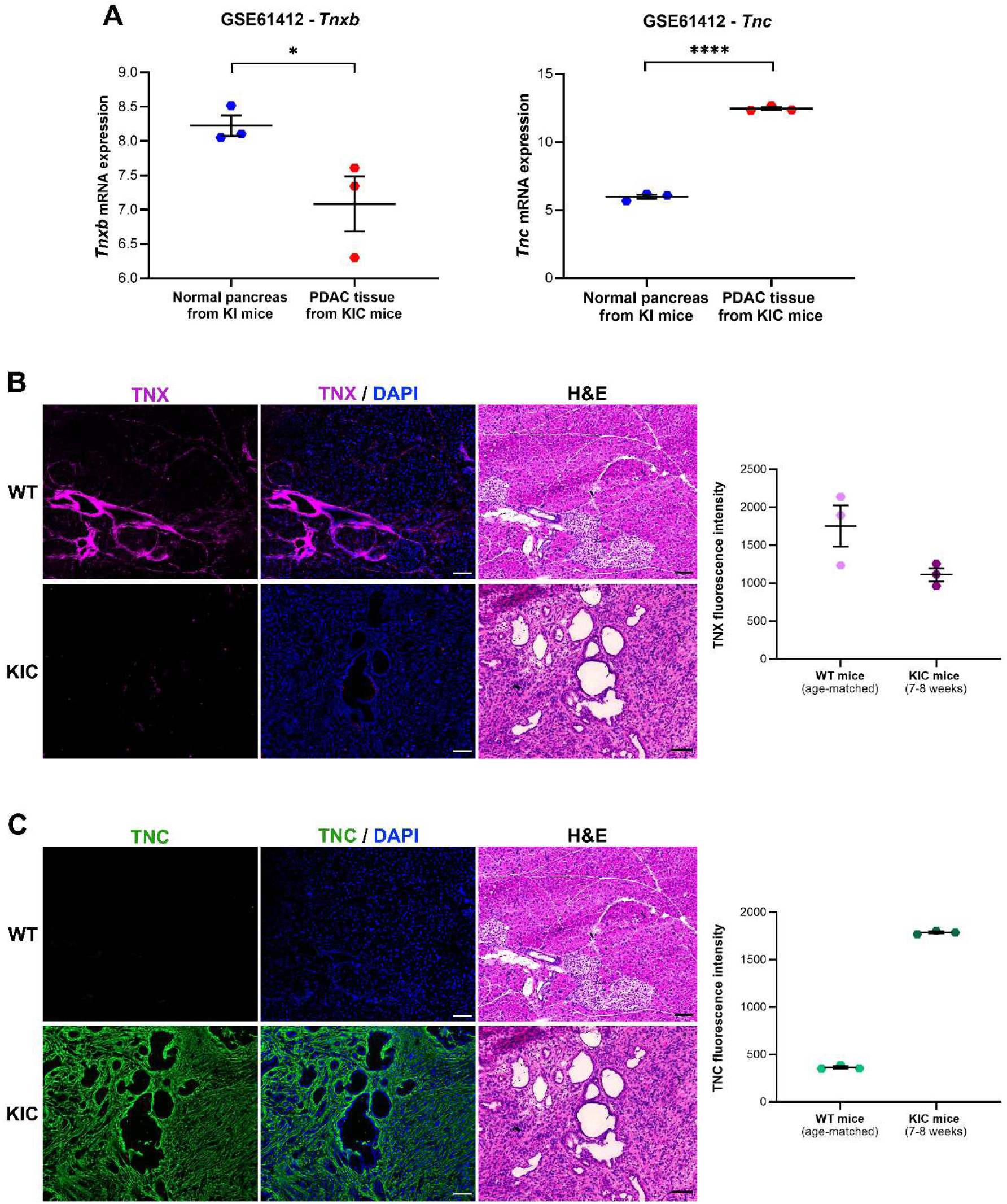
Tenascin-X mRNA and protein are lost in murine pancreatic tumors, in favour of Tenascin-C. **A**- Normalized *Tnxb (left)* and *Tnc (right)* expression values for the selected GEO dataset (individual values with mean ± SEM). **B**- Left: Representative pictures of pancreas from WT and KIC mice (7-8-week-old), labelled for TNX (magenta) and with DAPI (blue), with H&E staining performed on serial sections. Scale bar= 100 μm. Right: Quantification of TNX labelling intensity in the whole section of the pancreas of WT and KIC mice. **C**- Left: Representative images of pancreas from WT and KIC mice (7-8-week-old), labelled for TNC (green) and with DAPI (blue), with H&E staining performed on serial sections. Scale bar= 100 μm. Right: Quantification of TNC labelling intensity in the whole section of the pancreas of WT and KIC mice.

### TNX/TNC levels in mesenchymal cells are oppositely regulated by TGF-**β**

For more than 30 years, the literature has been regularly enriched by studies highlighting the role of TGF-β not only in the positive regulation of TNC in physiological (34,35) and pathological conditions (36–38), but also in PDAC progression (39). We thus hypothesized that the opposing regulation of TNX and TNC during pancreatic carcinogenesis could be attributable to TGF-β upregulation in Cancer-Associated Fibroblasts (CAFs). Therefore, we first confirmed in both previously selected microarray datasets, GSE62452 and GSE15471, that *TGFB1* mRNA levels were significantly increased in PDAC samples compared to NAT (Figure 4A) as already reported in previous studies (40). We then studied *in vitro* the effect of mesenchymal cell stimulation by TGF-β1 on TNX and TNC production. Experiments using Normal Human Dermal Fibroblasts (NHDF) and using SB431542 (a TGF-β receptor inhibitor) demonstrated that the absence of serum before and during TGF-β stimulation is required for cellular response (Supplementary Figure S2). Indeed, Fetal Bovine Serum (FBS) used in cell culture contains substantial doses of TGF-β, which bias cell stimulation by exogenous TGF-β (Supplementary Figure S2A and B). As expected, we showed by RT-qPCR and Western Blot on cell extracts, that TGF-β1 led to a significant increase in TNC levels. Very interestingly, TNX, which was highly produced by unstimulated NHDF, drastically disappeared following TGF-β1 cell stimulation (Supplementary Figure S2C and D). Opposite dysregulation of TNX and TNC levels by TGF-β1 was further confirmed *in vitro* on CAFs derived from [*Pdx1-Cre; LSL-Kras^G12D^; LSL-Trp53^R172H^*] (KPC) mice (Figure 4B) and on commercialized PSC (hPaSteC) extracted from the pancreas of a healthy patient (Figure 4C and D). It must be noted that other growth factors were also assayed for their capacity to modulate *Tnxb* and *Tnc* mRNA levels on KPC CAFs (Supplementary Figure S3). Thus, we demonstrated that cell stimulation by TGF-β3 also increased *Tnc* mRNA and decreased *Tnxb* mRNA, whereas FGF-2 decreased both *Tnc* and *Tnxb* mRNA levels. In our conditions, VEGF, BDNF and PDGF had no effect on TNC and TNX expression (Supplementary Figure S3).

**Figure 4:**
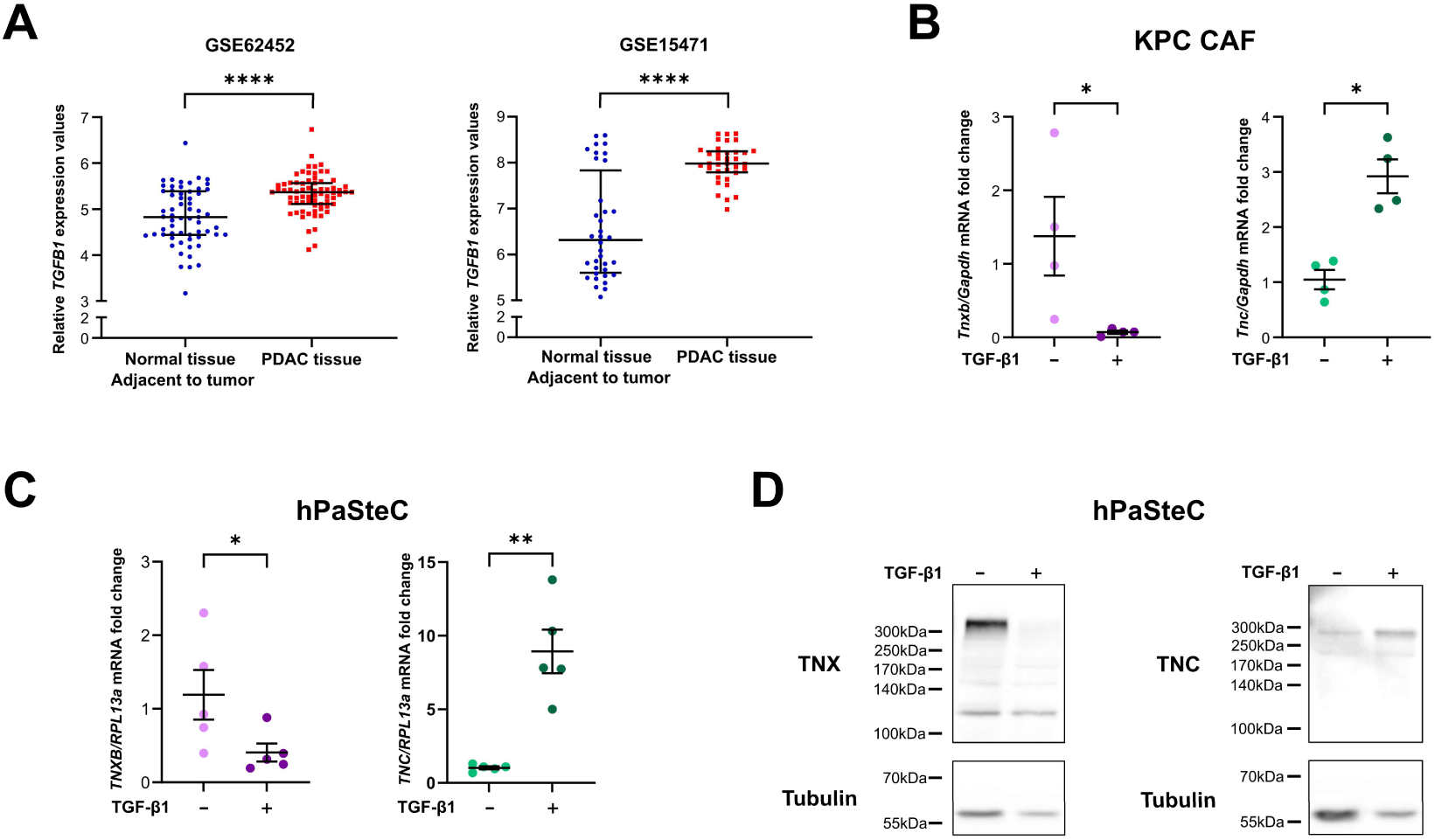
TGF-β1, which is over-expressed in PDAC, inhibits TNX production in vitro, in favour of TNC. **A**- Normalized *TGFB1* expression values for the two previously selected GEO datasets (individual values with mean± SEM). **B**- *Tnxb* (left) and *Tnc* (right) mRNA expressions in CAFs extracted from KPC mice following their stimulation with 20 ng/ml TGF-β1 for 24h. **C-** *TNXB* (left) and *TNC* (right) mRNA levels in normal human Pancreatic Stellate Cells (hPaSteC) following their stimulation with 10 ng/ml TGF-β1 for 24h. **D-** TNX (left) and TNC (right) protein production by hPaSteC following their stimulation with 10 ng/ml TGF-β1 for 48h. The presented western blots are representative of 4 independent experiments (see Supplementary Figure S4).

### Tenascin-X is transiently deposited during the initial steps of pancreatic carcinogenesis but not in reactive stroma

In order to determine more precisely when and how TNX disappeared during pancreatic carcinogenesis, we used the [*Pdx1-Cre; LSL-Kras^G12D^*] (KC) mouse model which is a less aggressive model of PDAC development than the KIC and KPC models. Pancreatic cancer development was followed by harvesting pancreas from 50-, 100-, 150- and 200-day-old KC mice. Acinar-to-ductal metaplasia (ADM) and PanIN-like lesions were delineated according to their morphological and molecular hallmarks (Figure 5A) and TNX positive areas around these lesions were analyzed (Figure 5B). PanINs were considered as low grade when epithelium presented a flat or papillary morphology without nuclear aberration, and as intermediate grade when nuclear polarity was at least partially lost and when cells entered into the duct. Finally, when epithelium was clearly pluri-stratified with many cells invading duct of the lesions, they were considered as high grade PanIN (Figure 5A). Cytokeratin-19 (CK19), a cytoskeletal protein of ductal cells, which is one of the main markers of ADM and thus of pancreatic pathogenesis, was used to confirm this evaluation. In normal pancreas, TNX was observed around ducts and vessels, and in very thin connective trabeculae around acini lobules (Figure 5B, top left). Surprisingly, we observed a biphasic TNX protein production during pancreatic carcinogenesis, with an increase during the early phases (ADM to intermediate grade PanIN) followed by a decrease, and ultimately a disappearance in high grade lesions, as observed in human samples (Figure 5B).

**Figure 5:**
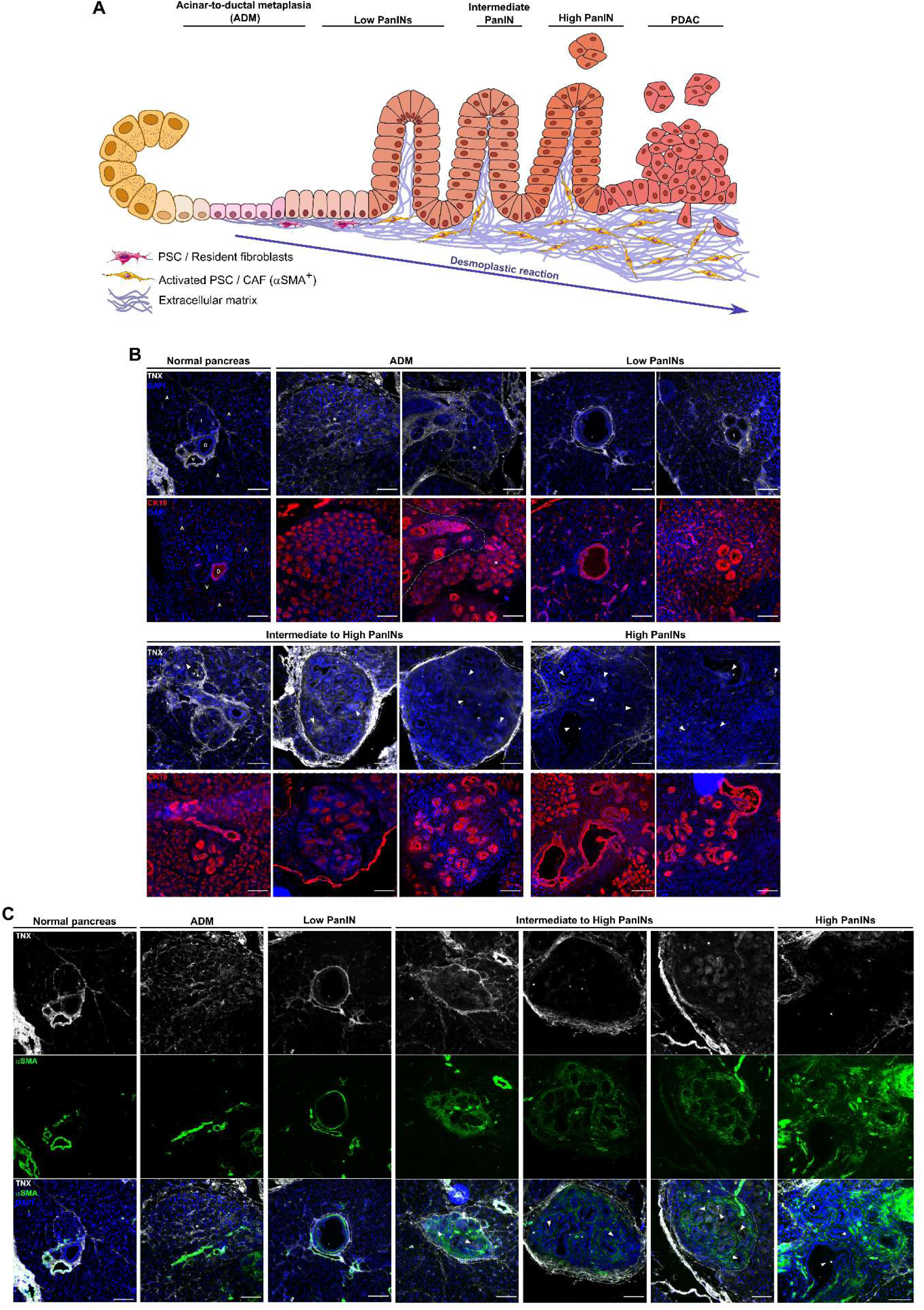
Tenascin-X is deposited in the initial steps of pancreatic carcinogenesis, but does not co-localized with reactive stroma. **A**- Schematic representation of linear model of PanIN progression, with associated desmoplastic reaction (reactive stroma development). **B**- Representative images of normal pancreas and the spectrum of PanIN-like lesions identified in KC mouse pancreas immunolabelled for TNX (white) and CK19 (red) on serial cryosections. On top left, normal pancreas is presented, with duct (D), vessel (V) and Langerhans islet (I) in the center, and acini (A) around. On top middle and right, ADM zones (*) and early PanINs are presented, respectively. Below, representative images of later stages PanINs are presented. **C**- Representative images of normal pancreas, ADM (*) and the spectrum of PanIN-like lesions identified in KC mouse pancreas immunolabelled for TNX (white, on top) and α-SMA (green, at the middle) on the same cryosections. The merged images are presented at the bottom. **B and C**- PanINs are surrounded by dotted line, and arrowheads highlight cells entering in duct, reflecting advanced PanINs. Some representative images in B are reused in C. Scale bar= 100 µm.

In order to determine more precisely at which stage TNX was involved in desmoplastic reaction accompanying PDAC, we performed a co-immunofluorescent staining of both TNX and α- SMA, a marker of activated fibroblasts involved in stroma reaction and associated with poor prognosis (41). In early lesions, TNX surrounded hyperplastic ducts and cells undergoing ADM, while α-SMA staining was quite weak and, when present, was closer to the lesions (Figure 5C, left). In advanced lesions, α-SMA staining accumulated around abnormal ducts whereas TNX faded around these ducts to form a dense sheath around the corresponding group of lesions (Figure 5C, right). The mutually exclusive α-SMA and TNX expressions are consistent with TGF-β production during pancreatic carcinogenesis because, unlike TNX, α-SMA expression is dependent on TGF-β production and is representative of tumor stroma (42). Taken together, these results demonstrate that before being downregulated, TNX is highly deposited around precursor lesions suggesting a reaction of the healthy stroma in response to lesion development.

### *TNXB* loss of expression in human PDAC is associated with increased proliferation

To decipher the consequences of TNX loss in human PDAC, and to get insights into the role of TNX in pancreatic cancer, we analyzed GSE62452 dataset containing the highest number of patients. We divided the cohort into *TNXB*^low^ and *TNXB*^high^ groups based on *TNXB* expression in PDAC samples. We then determined the differentially expressed genes by comparing *TNXB*^low^ and *TNXB*^high^ groups with GEO2R (Supplementary Table S5), and performed enrichment analyses using the computational tools Metascape (https://metascape.org/) (43) and Reactome Pathway Knowledgebase (https://reactome.org/) (44) (analyses carried out on Nov. 2024). Through Metascape analysis, we clearly established that in *TNXB*^low^ tumors, the top 20 enriched terms (-log10(*P*)>10) were linked to cell division, cell cycle checkpoint and DNA repair (11 terms), morphogenesis and embryogenesis (4 terms) and cancer, ECM, adhesion and signalling for the other terms (Figure 6A). The Reactome analysis tool confirmed these results and enabled to identify multiple genes over-represented in *TNXB^low^* group (*CDK1, CCNB1, TOP2A, CDC6, CDH1, FOXM1*, *etc.*). Interestingly, those genes are linked to PDAC progression, prognosis and/or patient survival and considered as targets for PDAC treatment (45–48) (Figure 6B and Supplementary Table S5). Metascape and Reactome enrichment analyses were also conducted using the differentially expressed genes in *TNC*^high^ and *TNC^low^* groups with GEO2R (Supplementary Table S6). As expected, the top 20 enriched terms (- log10(*P*)>10) in *TNC^high^* group were mainly linked to ECM, cell adhesion and migration (8 terms). Among the differentially expressed genes in both groups, we noticed collagen genes and genes involved in collagen processing (*BMP1, COL1A1, COL1A2, COL3A1, COL4A1*, *etc.*) and ECM remodelling (*ADAM12, ADAM17, ADAM19, ADAMTS6, ADAMTSL1, SERPINE1, SERPINH1*, *etc.*). Other terms were significantly enriched such as smooth muscle contraction and cytoskeleton (2 terms), development (5 terms) and cell signalling (Figure 6C and D). Among the genes annotated with smooth muscle contraction, we noticed *ACTA2, MYH11* and *MYL9*, 3 markers of myofibroblastic CAFs (myCAFs) which are adjacent to neoplastic cells and produce desmoplastic stroma (49). Interestingly, *TPM2* and *TPM4*, which encode cytoskeleton proteins, are over-expressed in *TNC^high^* tumors. Unlike *TPM1* and *TPM2* which are directly linked to PDAC and expressed by myCAFs upon TGF-β stimulation, *TPM4* has never been described in PDAC (49). Furthermore, by comparing enriched terms in *TNXB^low^ versus TNXB^high^* tumors and *TNC^high^ versus TNC^low^*tumors, we observed that the 2 Tenascin family members mostly impacted different biological processes (Figure 6B and D). Altogether, these results suggest that (i) TNX could act during PDAC progression by inhibiting cell proliferation/division and that (ii) TNC would be linked to ECM remodelling and stiffness.

**Figure 6:**
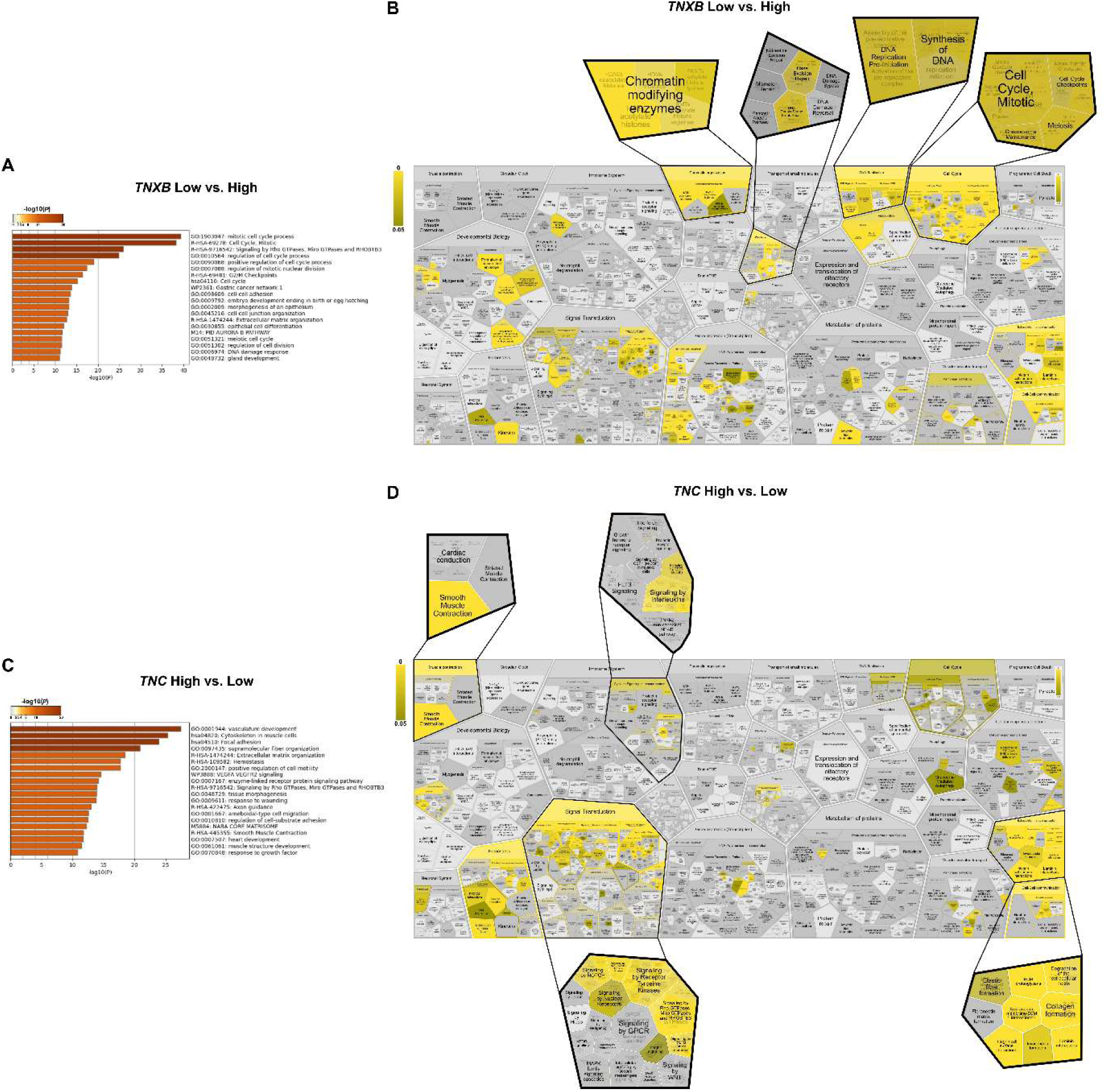
Low TNXB expression in PDAC samples is correlated with cell proliferation, as demonstrated by bioinformatic analysis of the GSE62452 dataset. **A**- Biological Processes enriched in the upregulated genes identified in the *TNXB^low^* PDAC samples, determined by Metascape tool and ranked according to their p-value. **B-** Flattened representation of enriched biological pathways identified by Reactome in over-expressed genes in TNXB^low^ tumors. **C-** Biological Process enriched in the upregulated genes identified in the *TNC^high^* PDAC samples, determined by Metascape tool and ranked according to p-value. **D-** Flattened representation of enriched biological pathways identified by Reactome in over-expressed genes in TNC^high^ tumors. **A and C-** The color scale ranges from brown (the smallest p-value) to light yellow. **B** and **D-** Homo sapiens pathways represented as proportionately packed cells, each cell representing a Reactome pathway. The cell areas are proportional to the number of proteins annotated in the pathway and its sub-pathways in humans (https://www.ebi.ac.uk/training/online/courses/reactome-exploring-biological-pathways/understanding-the-pathway-browser/diagram-panel/pathway-overview/). The color scale ranges from yellow (the smallest p-value) to dark.

To validate TNX role in cell proliferation, we analyzed, by immunohistochemistry, TNX, TNC and Ki67 localization on serial sections of human PDAC samples (Figure 7A). Considering that TNX expression is significantly downregulated in advanced lesions and in PDAC, we focused on PanINs lesions that can be found in tumor samples. We could clearly identify lesions surrounded by strong TNX deposition with no TNC and in which the proliferation of epithelial cells (as determined by Ki67) was almost undetected. On the contrary, in other lesions where TNX was absent, TNC was heavily deposited around these lesions, and cells showed strong expression of Ki67. We classified the lesions according to their negative or positive status for TNX or TNC and determined the percentage of Ki67-positive cells within these lesions (Figures 7B and C). Two graphical representations are shown: on the left, each patient has been analyzed individually and, on the right, each lesion of each patient is considered separately (Figures 7B and C). This last representation takes into account the potentially different evolution of each lesion. There was a lower percentage of Ki67-positive cells in TNX-positive lesions and a significantly higher percentage of Ki67-positive cells in TNC-positive lesions (Figure 7B and C). These findings were further confirmed in the aggressive KIC mouse model with regard to tumor cell proliferation (Figure 7D-G). Indeed, we observed in the connective tissue of WT mouse pancreas, high levels of TNX and low levels of TNC correlating with low numbers of Ki67-positive cells. In contrast, in pancreatic samples from KIC mice, TNX was hardly detectable compared to TNC, and this was associated with an increase in proliferating (Ki67^+^) cells (Figure 7D-G). These results highlight the crucial role of the TNX/TNC ratio during PDAC development which could be mediated at least in part by its impact on tumor cell proliferation.

**Figure 7:**
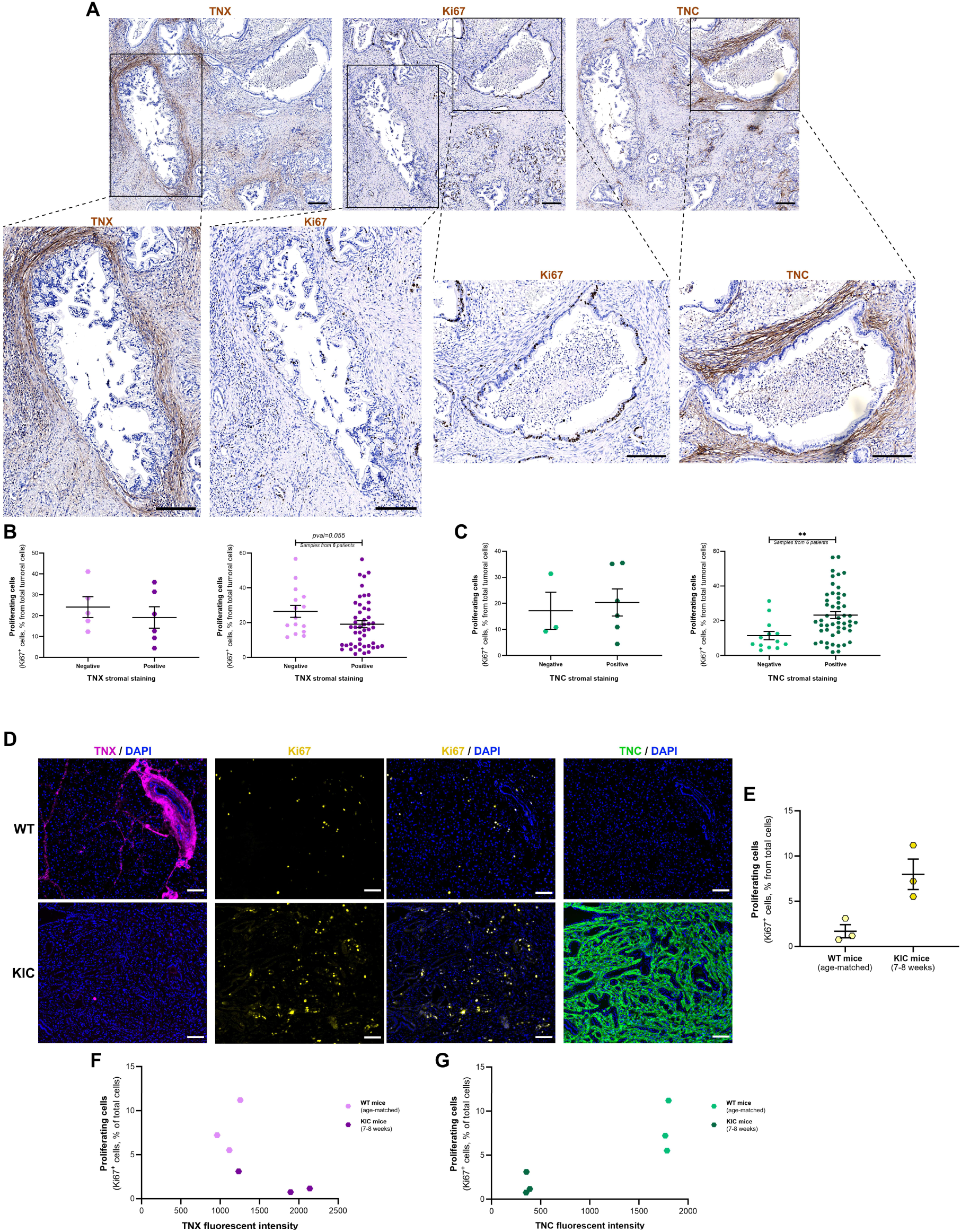
Tenascin-X and -C depositions are respectively associated with low and high proliferation in human and GEMM samples. **A**- Representative images of human PDAC samples immunolabelled for TNX, TNC and Ki67 on serial sections (top). Higher magnification of the images presented at the top showing one tumor gland weakly stained for Ki67 and surrounded by TNX but not TNC, while the other is rich in Ki67^+^ cells and is surrounded by TNC bot not TNX (bottom). Scale bar= 200 µm. **B**- Quantification of proliferating cells (Ki67^+^ cells) depending on stromal TNX. The Left graph shows the mean of the different regions analyzed for the 6 patients and the right one all the regions of interest analyzed (all the patients). **C**- Quantification of proliferating cells (Ki67^+^ cells) depending on stromal TNC. The left graph shows the mean of the different regions analyzed for the 6 patients, and the right one all the regions analyzed (all the patients). **D**- Representative images of WT and KIC mouse pancreas (7- to 8-week-old), labelled for TNX (magenta), Ki67 (yellow), TNC (green) and with DAPI (blue). Scale bar= 100 µm. **E**- Quantification of proliferating cells (Ki67^+^ cells) in WT and KIC mouse pancreas. Between 54 831 and 198 074 cells per mouse were counted. **F**- Correlation between the % of proliferating cells and TNX intensity in the same pancreas (one point per mouse). **G**- Correlation between the % of proliferating cells and TNC intensity in the same pancreas (one point per mouse).

## Discussion

We demonstrated in this study that *TNXB* gene expression and TNX protein level were drastically decreased in human PDAC tumors compared to normal tissue adjacent to the tumor or to normal pancreas samples, and that low *TNXB* expression seemed to be correlated with poorly differentiated tumors (Figure 1H), which are often more aggressive, and was associated with reduced patient survival (Figure 1I). These results are consistent with our previous results showing that *TNXB* transcript and TNX protein levels in tissue are decreased in 13 cancer types including the 6 most prevalent cancers worldwide (29). Thus, the down-regulation of TNX expression seems to occur in most solid cancers. In contrast to TNX, TNC was upregulated at the transcriptional and translational levels in PDAC compared to NAT. However, a correlation between high TNC expression and disease aggressiveness or patient survival was not overt. These results are consistent with previous studies demonstrating that TNC gradually increases from low grade PanIN lesions to pancreatic cancer (8,50–52), and that TNC does not appear to be a prognostic marker of PDAC, except when present in specific area such as perineural sites at the invasive tumor front (8,53).

Here, we also demonstrated that opposite dysregulation of TNX and TNC may be attributable, at least partially, to TGF-β1 growth factor presence (Figure 4). Interestingly, all members of the TGF-β family were upregulated in PDAC compared to NAT samples in the 2 selected GSE datasets (GSE15471 and GSE62452) (Supplementary Figure S7). TGF-β is known to play pleiotropic roles in cancer progression by (i) promoting ADM, (ii) enhancing cancer cell invasion, dissemination and stem cell properties, (iii) shaping the TME through activation of various cell types and notably CAFs, and (iv) repressing the anti-tumor functions of various immune cell populations (39,54,55). TGF-β blocking antibodies have been developed for PDAC treatment and showed promising results in orthotopic models of pancreatic cancer by enhancing sensitivity to combination chemotherapy (56,57). Interestingly, in this latest study, single-cell transcriptomic analysis of fibroblasts extracted from orthotopic tumors was performed and various differentially expressed genes were dysregulated following TGF-β blocking antibodies treatment. Among them, *TNXB* and *TNC* genes featured prominently, with increased expression of *TNXB* and decreased expression of *TNC* in the inflammatory CAF (iCAF) sub-population, which is known to be associated with prolonged survival (57). In combination with our study, these results reinforce the role of TGF-β in Tenascin family member regulation and in TME remodeling and encourages ongoing efforts to target TGF-β and its signaling pathways for PDAC treatment.

Changes in TNX and TNC during pancreatic carcinogenesis were confirmed in the aggressive KIC mouse model and led us to characterize the kinetics of TNX loss during pancreatic carcinogenesis using the KC mouse model. Whereas we were expecting a progressive decrease of TNX deposition during pancreatic carcinogenesis, we showed that TNX presence was first reinforced around cells in ADM and around low-to-intermediate grade PanINs, before being drastically reduced or lost in high grade PanINs. To better understand TNX deposition at the initial stages of PanIN progression, we assessed whether TNX was associated with stromal reaction and thus produced by the CAFs (Figure 5C). CAFs are key components of the pancreatic tumor microenvironment, maintaining the extracellular matrix, while also being involved in intricate crosstalk with cancer cells and infiltrating immunocytes (58). The most commonly used marker to detect CAFs is α-SMA, although CAFs are very heterogeneous and this marker is more specific of CAFs with contractile behaviour, *i.e.* myCAFs (41,58). We clearly showed that TNX deposition did not co-localize with α-SMA staining, thus suggesting that TNX might not be produced to favor PDAC progression but in contrary to counteract pre-tumoral lesion development during the first steps of pancreatic carcinogenesis. Analysis of single-cell transcriptomic datasets using TISCH2 software (http://tisch.comp-genomics.org/home/) indicates that iCAFs might express *TNXB* genes as recently suggested by Mucciolo and colleagues (59). It has been recently proposed that the modification in microenvironment could precede pancreatic (pre)tumoral cell development (60), and that the desmoplastic reaction can be considered as a physical barrier against tumor cell invasion. In line with this hypothesis, the development of a protective ECM has been demonstrated in first steps of tumor progression in other tumor types (61). Thus, ECM microenvironment could be spatially and temporally compartmentalized during carcinogenesis depending on cancer cell influence and plays a dual role, anti-tumoral at the early stage and pro-tumoral at the late stage. The peripheral localization of TNX during pancreatic carcinogenesis and the strong deposition of TNC in some specific human PDAC lesions lacking TNX, could reflect these two compartments. Furthermore, it has been shown that TNC is deposited in close proximity to tumor cells, but not in peripheral stroma (62). Other ECM molecules are heterogeneously deposited within PDAC stroma. The identification of the proteins deposited concomitantly with TNX in the peripheral sheath surrounding lesions would allow a better understanding of the role of perilesional ECM and of TNX in the first steps of carcinogenesis, and is a prerequisite to design efficient therapies. These investigations could be done in GEMMs mimicking PDAC development and in human pancreatic samples with various stages of PanINs although the late diagnosis of PDAC makes the collection of such samples challenging.

Our work also highlights the importance of histologically investigating ECM in PDAC to better characterize the different types of deposited matrix. Most previous studies describing the matrix accompanying carcinogenesis use whole tissue analysis (63–65), which does not permit the discrimination of ECM heterogeneity within the stroma of early pancreatic lesions. Our work, showing two lesions in close proximity with two different TN deposits, raises the following questions: what distinguishes those lesions and which cells produce this matrix? Two main PDAC subtypes have been identified mainly by transcriptomic and histological analyses of the epithelial compartment: a subtype more frequently resectable, with a higher level of differentiation and often associated with fibrosis and inflammation, and a basal-like subtype with a poorer clinical outcome and loss of differentiation (66). However, the in-depth analysis of the morphology of these lesions by anatomical pathologists specialized in pancreatic cancer did not allow us to distinguish them.

CAFs promote tumor progression by depositing a dense ECM around lesions. However, the limited success of therapeutic approaches aimed at CAF depletion suggests that some elements of CAF behaviour could be cancer-restraining (rCAFs) during the earliest stages (11,67). rCAF populations have been recently characterized (68,69), which could corroborate our idea of a spatio-temporally compartmentalized stroma during PDAC progression. In addition, different types of stroma have been characterized and linked to different clinical outcomes, depending on their composition (collagen content, CAF markers, *etc*) (70), or their position relative to tumor cells (62). Therefore, TNX could be expressed by resident fibroblasts or particular CAFs in normal pancreas and in the initial steps of pancreatic cancer and TNC could be expressed by pro-tumoral CAFs in later stages of pancreatic carcinogenesis possibly under the influence of soluble factors released by tumor cells such as TNF-α and TGF-β (50,52).

The biological processes enriched in *TNXB^low^* tumors (Figure 6A and B) and in *TNC^high^* tumors (Figure 6C and D) were identified using Metascape and Reactome Pathway Knowledgebase. Many genes involved in cell cycle, and notably in cell proliferation, such as cyclins, histones, mini-chromosomal maintenance and centromere-associated genes, were upregulated in *TNXB^low^* tumors and downregulated in *TNC^high^* tumors, indicating that TNX could inhibit tumor and/or stromal cell proliferation and TNC could favor cell cycle progression. To confirm this hypothesis, various experiments were conducted on human PDAC samples and PDAC mouse model and we demonstrated that the TNX/TNC balance was correlated to cell proliferation. Those results are in agreement with previous studies performed on pancreatic cancer cell lines and human PDAC samples showing that TNC promotes pancreatic cancer cell growth *via* integrin pathway activation (51,71). Interestingly, the role of TNX as a proliferation inhibitor echoes one of its referencing names (growth-inhibiting Protein 45) by Kim and colleagues in 2007.

In conclusion, we report that TNX is deposited in the first steps of carcinogenesis, but, unlike TNC, is no longer expressed at a detectable level in PDAC. The TNX/TNC ratio is at least modulated by TGF-β family members and its variations are correlated with tumor and/or stromal cell behavior modifications, in particular tumor cell proliferation. The underlying mechanism is still unknown but currently under investigation in our laboratory and could lead to the emergence of TN family as potential stromal targets in PDAC treatment.

## Materials and Methods

### *In vitro* experiments

#### Cell Culture

Normal Human Dermal Fibroblasts (NHDF) were isolated from abdominal adult skin explants obtained from the Biological Resource Center GCS/CTC (AC-2019-3476, 2^nd^ July 2020) hosted by Hospices Civils de Lyon (France). NHDF were cultured in DMEM/F-12 (Gibco) supplemented with 10% FBS and 1% Pen-Strep. Human Pancreatic Stellate Cells (hPaSteC), extracted from normal pancreas, were purchased from ScienCell Research Laboratories and were cultured in Stellate Cell Medium (SteCM) according to the supplier’s protocol. Murine CAFs were extracted from [*Pdx1-Cre; LSL-Kras^G12D^; LSL-Trp53^R172H^*] (KPC) mice (according to previously published protocol, (72,73)) and were cultured in DMEM (Gibco) supplemented with 10% FBS and 1% Pen-Strep. NHDF and murine CAFs were cultured for no more than 20 passages at 37°C with 5% CO_2_ and hPaSteC for no more than 5 passages. Mycoplasma testing of cells was performed routinely prior to freezing. To analyze the production of Tenascins, cells were first serum-starved (for 24h for NHDF and 16h for hPaSteC) and then stimulated or not with 10-20 ng/ml TGF-β1 (PeproTech®), 20 ng/ml TGF-β3, FGF-2, EGF, VEGF, BDNF or PDGF-AA (Sigma-Aldrich) in serum-free medium for 24h (for mRNA) to 72h (for protein). Cells were then collected for mRNA or protein analyses.

#### RNA extraction and quantitative real time PCR

Total RNA was isolated using RNeasy Plus Mini kit according to manufacturer’s instructions (Qiagen). Reverse transcription was performed with 500ng RNA following the protocol of the PrimeScript RT reagent kit (Takara). The cDNA was treated with 1 U of Escherichia coli RNase H (Roche). qPCR was performed using FastStart Universal SYBR Green Master mix (Roche) in a Rotor-Gene Q system (Qiagen) with specific primers (Supplementary Table S7). Results were expressed as relative values normalized with the reference gene RPL13A and quantified by the 2−ΔΔCt method. The mean of ΔCT values for unstimulated cells was set at 1, and expression data for each sample are presented relative to this reference value.

#### Protein extraction and Western blot analyses

Proteins were extracted using the following lysis buffer: 150 mM NaCl, 50 mM Tris pH 8 and 1% (v/v) NP-40 supplemented with complete protease inhibitor and phosphatase inhibitor cocktails (Roche). Protein concentration was determined using a BCA assay (ThermoFisher Scientific). For Western blot analyses, samples were separated by SDS-PAGE and blotted onto PVDF membranes. Specific antibodies were used: anti-TNX (polyclonal, KR86 or KR87 kindly provided by Dr. Manuel KOCH, recognizing the C-terminal end of the full-length human protein, 1/500), anti-TNC (monoclonal, MAB2138, R&D Systems, raised against mouse immature astrocyte-derived Tenascin C, 1/250), anti-acetylated Tubulin (monoclonal, T7451, Sigma-Aldrich, 1/1000). The absence of cross-reactivity between anti-TNX and anti-TNC antibodies was verified. HRP-coupled secondary antibodies were used with a chemiluminescent kit (Amersham or ThermoFisher Scientific) to detect the immunoreactive proteins with a FX Fusion CCD camera (Vilber). Quantitative analyses were performed using the Fiji software.

### Human tissue analyses

#### In silico *analyses*

To study *TNXB* and *TNC* gene expressions in PDAC tumors from patients compared to “healthy” pancreas, microarray datasets were selected from the NCBI GEO data repository (https://www.ncbi.nlm.nih.gov/gds). Two datasets (GSE62452 and GSE15471, 60 and 36 patients, respectively) were analyzed with GEO2R, an analysis tool which compares 2 groups using LiMMA (Linear Models for Microarray Analysis) R package (Bayesian statistics). Differentially expressed genes (DEGs) were extracted (Benjamini & Hochberg False Discovery Rate adjusted *p*-value<0.05), and the first probes/identifiers (ID) detected in the list, corresponding to *TNXB* and *TNC* genes, were used to plot and compare their expression values in PDAC *versus* NAT using paired t-test after checking the Gaussian distribution. The same tool was used to analyze the GSE61412 dataset and to compare *Tnxb* and *Tnc* expressions in PDAC from KIC mice compared to equivalent samples from KI mice (mice devoid of the Cre recombinase construct and thus considered as WT mice) as control.

Patient survival and enriched pathways were studied on the GSE62452 dataset. First and last quartiles based on *TN* expression values in tumor samples allowed to define 2 groups: *TN*^low^ and *TN*^high^ tumors, respectively (25% of patients with the lowest values = *TN*^low^, 25% with the highest values = *TN*^high^). Cumulative survival proportions were estimated using the Kaplan– Meier method and *TN*^low^ and *TN*^high^ groups were compared thanks to Log-Rank (Mantel-Cox) statistical test, and GEO2R. Computational enrichment analyses of DEGs were performed on November 2024, using Metascape, which provides provide a comprehensive gene list annotation and analysis resource for experimental biologists (43), and the Reactome Pathway Knowledgebase, which is focused on biological pathways (44).

#### Immunohistochemistry on human tissue samples

For TN immunodetection on human PDAC tissue, commercially available Tissue Micro Array (TMA) slides - including 62 tumors and matched normal tissues adjacent to the tumor (NAT) (US Biomax, HPanA150CS03) - were used. Immunohistochemistry (IHC) staining was performed using R.T.U. Vectastain Universal Elite ABC Kit (VectorLabs, PK-7200) following manufacturer’s instructions. After deparaffinization and rehydration, antigen retrieval was achieved in sodium citrate buffer (pH 6) for 20 min at 98°C followed by a 20 min cooling time. Then, endogenous peroxidases were quenched by 3% (v/v) H_2_O_2_ and sections were saturated with normal horse serum. Primary monoclonal antibodies (anti-TNX: sc-271594, Santa Cruz Biotechnology, recognizing the 14 amino-acid from the C-terminal end of the human protein, 1/100, anti-TNC: sc-59884, Santa Cruz, raised against purified full-length native Tenascin-C of human origin, 1/2000 or anti-Ki67: GTX16667, GeneTex, 1/500) were incubated in blocking solution overnight at 4°C and biotinylated-secondary antibody for 30 min at room temperature. Development was performed using ABC Reagent (30 min incubation) and 3,3′- Diaminobenzidine (DAB, VectorLabs, SK-4105). Finally, nuclei were counterstained with Gill’s Hematoxylin, sections were mounted with DEPEX and tissue cores were scanned using AxioScan.Z1 microscope (Zeiss).

#### Image analysis

Each imaged core was extracted by QuPath software (https://qupath.github.io/) and analyzed using Fiji software (https://fiji.sc/) as previously described (29). Stromal and epithelial tissues were separated thanks to Fiji Trainable Weka Segmentation plugin (https://imagej.net/Trainable_Weka_Segmentation). The segmentation training was newly performed for each tissue core because of histological differences between tumoral and normal samples. Colors were then separated by “H DAB” Colour Deconvolution and DAB staining was quantified in stromal region (mean grey value). Reciprocal mean grey value (255 – mean grey value) was considered, as LookUp Table (LUT) is inverted in Fiji software. Additionally, TN protein staining intensity in stroma was blindly evaluated by an anatomical pathologist specialized in pancreatic cancers (Dr. Valérie Hervieu, Lyon, France (74,75)).

### Mouse tissue analyses

#### Animals

LSL-Kras^G12D^ (B6.129S4-*Kras^tm4Tyj^*/J strain, Jackson Laboratory, (76)) and *Pdx1*-Cre (B6.FVB-Tg(Pdx1-cre)6Tuv/J strain, Jackson Laboratory, (77)) mouse strains were previously described and respectively provided by Dr. Philippe Bertolino and Dr. Laurent Bartholin (CRCL, Lyon, France). Mice were maintained in a pathogen-free animal facility at the “ALECS SPF module” (Lyon, France) and crossed in order to obtain [*Pdx1-Cre; LSL-Kras^G12D^*] mice, also named “KC” mice (77). The KC mouse model presents the classical initiating mutation of *Kras* found in almost 95% of PDAC patients and leading to the constitutive activation of the oncogene *Kras.* The Cre recombinase expression is driven by the *Pdx1* promoter which is specifically expressed in early pancreatic progenitor cells. These mice are described to develop ductal lesions similar to human PanINs and at low frequency, these lesions progress to invasive and metastatic PDACs within one year (77). Animals were genotyped one week after birth and euthanized at 50-, 100-, 150- and 200-days post-parturition. The [*Pdx1-Cre; LSL-Kras^G12D^; Ink4a/Arf^lox/lox^*] (KIC) mouse model provided by Dr. Ana Hennino (CRCL, Lyon, France) presents (i) the classical initiating mutation of *Kras* and (ii) the mutation of the cyclin-dependent kinase inhibitor 2A (*Cdkn2a*) gene, found in 98% of PDAC, and leading to the loss of function of 2 different tumor suppressors INK4A and ARF (78). Thus, in this model, *KRAS* and *CDKN2A* mutations are specifically found in the pancreas and lead to the rapid progression of PanIN to metastatic PDAC within 11 weeks of age (79). Pancreases from 7-8-week-old KIC mice were collected at humane end-point and compared with equivalent tissue from age-matched wild-type (WT) mice. After dissection, pancreases were directly embedded in OCT (Optimal Cutting Temperature) compound, snap-frozen in cold isopentane and kept at −80°C until sectioning. All animal experiments were executed in compliance with institutional guidelines and regulations and after approval by an ethic committee under regulatory of governmental authority (APAFiS #16242-201804271428498 and #35125-2022020214089191).

#### Immunofluorescence

To analyze TNX and TNC levels in pancreas of KIC and KC mice, and their correlations with reactive stroma, immunofluorescence experiments were achieved on 10 µm cryosections. Immediately after sectioning, slices were fixed in cold methanol (−20°C) for 10 min and immunofluorescent staining was performed as follows. After a 10 min permeabilization step with 0.5% (v/v) Triton-X100 in Phosphate Buffered Saline (PBS), non-specific sites were blocked using normal horse serum (2.5 to 5% (v/v) in PBS). Primary antibodies (monoclonal anti-TNX, FN9/10, homemade antibodies raised against the FN-III_9-10_ domains of the bovine protein), 1/100 (21); monoclonal anti-TNC, MAB2138 (R&D Systems), 1/250; monoclonal anti-CK19, TROMA-III-s (DHSB), 1/100; polyclonal anti-α-SMA, GTX100034 (GeneTex), 1/2 000) were applied overnight at 4°C and secondary antibodies conjugated to a fluorochrome were incubated for 45 min at room temperature. Slides were then mounted using Fluoromount Medium with DAPI (VectorLabs, H-1200). In parallel and when necessary, Hematoxylin and Eosin staining was performed on serial sections in order to better identify pancreatic lesions.

#### Image analysis

All sections were imaged using the slide scanner AxioScanSP5 X (Zeiss) at CIQLE (Lyon, France). Images were quantified using QuPath (https://qupath.github.io/) software. A simple tissue detection was first made in order to measure the total area of pancreas in the image, and TNX staining was quantified thanks to Pixel classification in whole pancreas and around each identified lesion, using the same Brightness and Contrast values between sections.

### Statistical analyses

Statistical analyses were performed using GraphPad Prism 8 software (La Jolla, California, USA). Unless otherwise noted, data were presented as mean ± SEM. After image analysis, groups were compared either by non-parametric one-way ANOVA (Kruskal Wallis test, if more than 2 groups were compared) or non-parametric mean comparison tests (Wilcoxon Rank sum for paired data, Mann-Whitney for unpaired data). *P-*values<0.05 were considered significant (**** p<0.0001; *** p<0.001; ** p<0.01 and * p<0.05).

## Data Availability

The data analyzed in this study were obtained from Gene Expression Omnibus (GEO) at GSE62452, GSE15471 and GSE124731.

## Acknowledgments

We acknowledge the contribution of SFR Santé Lyon-Est (UAR3453 CNRS, US7 Inserm, UCBL1) through their animal facility (ALECS module SPF) and the microscopy platform (CIQLE: a LyMIC member) and particularly Bruno CHAPUIS for its precious help in the use of the slide scanner Zeiss Axio Scan.Z1. Additionally, we greatly thank members of the histology and imaging facility (PrImaTiss) at the LBTI institute. Finally, we acknowledge Dr. Philippe BERTOLINO and Dr. Laurent BARTHOLIN for providing us the Kras^G12D^-driven mouse model and for constructive advice regarding the project.

## CRediT author statement

**Sophie Liot:** Conceptualization, Methodology, Data Curation, Software, Investigation, Formal Analysis, Visualization, Writing – Original draft; **Céline Schmitter:** Investigation; Visualization, Data curation, Formal Analysis, Writing – Original draft; **Perrine Mercier-Gouy:** Investigation, Validation; **Naïma El Kholti:** Methodology; **Jonathan Balas:** Investigation; **Laura Prigent:** Validation; **Alexandre Aubert:** Validation; **Claire Lethias:** Resources; **Jessica L. Chitty:** Supervision, Funding Acquisition, Writing - Review and Editing; **Manuel Koch:** Resources; **Tanguy Fenouil:** Resources, Data Curation; **Thomas R. Cox:** Resources, Supervision, Funding Acquisition, Writing - Review and Editing; **Ana Hennino:** Resources; **Valérie Hervieu:** Resources, Investigation, Data curation; **Sylvie Ricard-Blum:** Software, Investigation, Formal Analysis, Writing - Review and Editing; **Ulrich Valcourt:** Funding Acquisition, Supervision, Writing - Review and Editing; **Elise Lambert:** Conceptualization, Methodology, Visualization, Supervision, Writing – Original draft, Funding Acquisition.

## Data sharing statement

The data that support the findings of this study are available upon request from the corresponding author.

