## Supplementary Figures for "Dynamic dysregulation of Tenascin-X/Tenascin-C balance controlled by Transforming Growth Factor-β leads to tumor cell proliferation during pancreatic carcinogenesis"

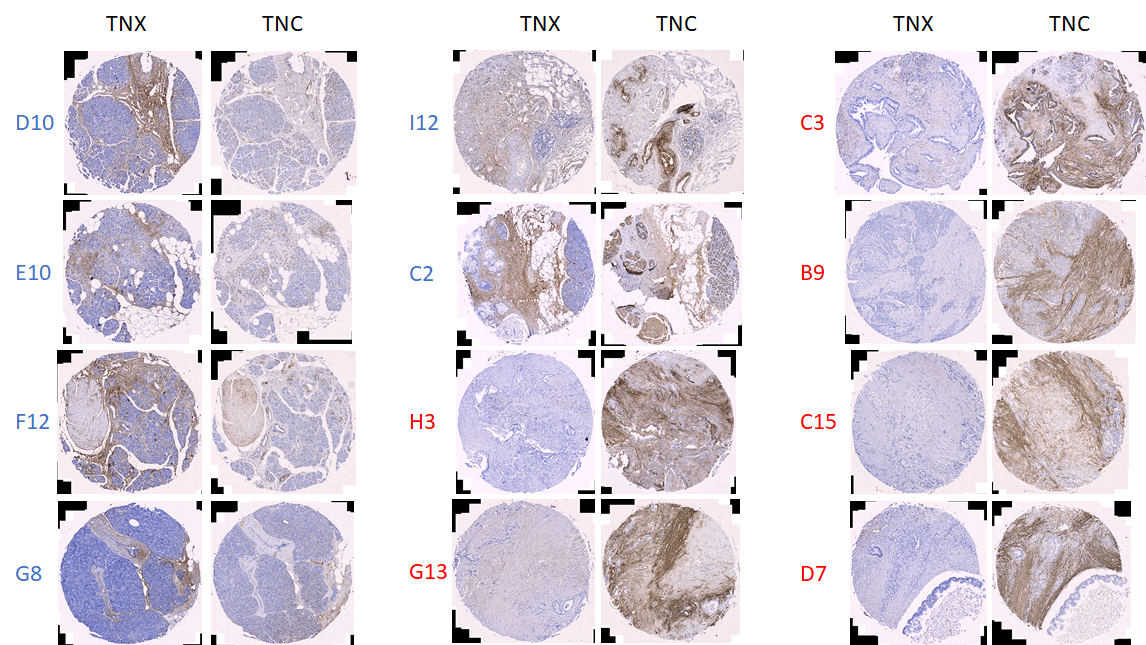

***Supplementary Figure S1: Opposite deposition of TNX and TNC in NAT and tumor pancreatic samples.*** *Serial slices of TMA were immunolabelled with anti-TNX or -TNC antibodies. Representative cores were selected (6 for NAT and 6 for tumor samples). Core position is indicated on the left of the 2 labelling and is in blue for NAT samples or red for tumor samples.*

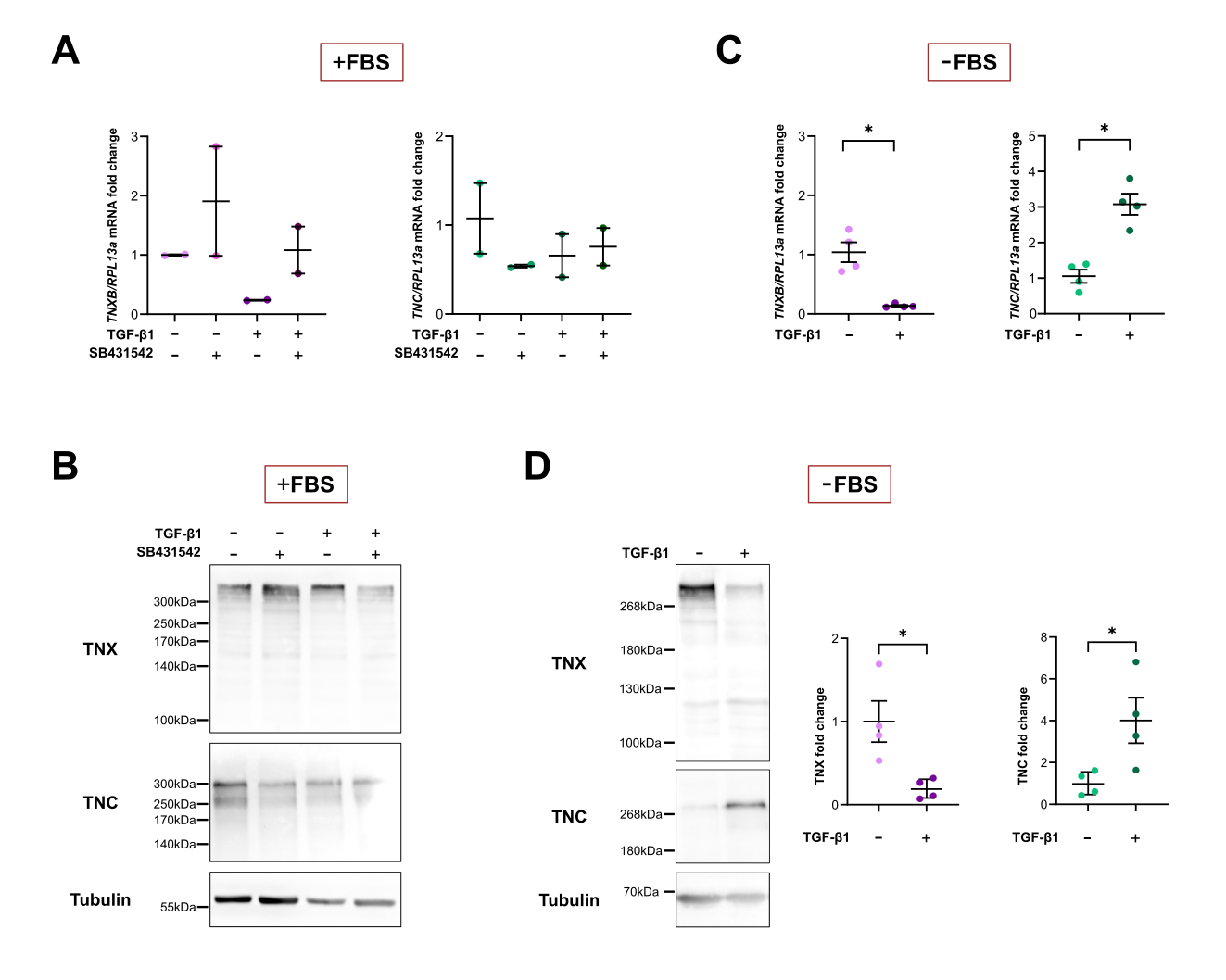
 ***Supplementary Figure S2: Regulation of TNX and TNC productions in Normal Human Dermal Fibroblasts (NHDF) following their stimulation with TGF-β1.*** *NHDF were serum-starved (-FBS, C and D) or not (+FBS, A and B) for 24h and then stimulated or not with 10 ng/ml TGF-β1 for 48-72h in presence or not of 10µM SB431542 (a selective inhibitor of TGF-β Type I Receptor/ALK5). mRNA or protein extracts were analyzed by qRT-PCR (A and C) or Western Blot (B and D), respectively. The Western Blot presented in D (left) is representative of 4 independent experiments and TNX and TNC productions have been quantified through Fiji analysis and normalized to acetylated Tubulin (right).*

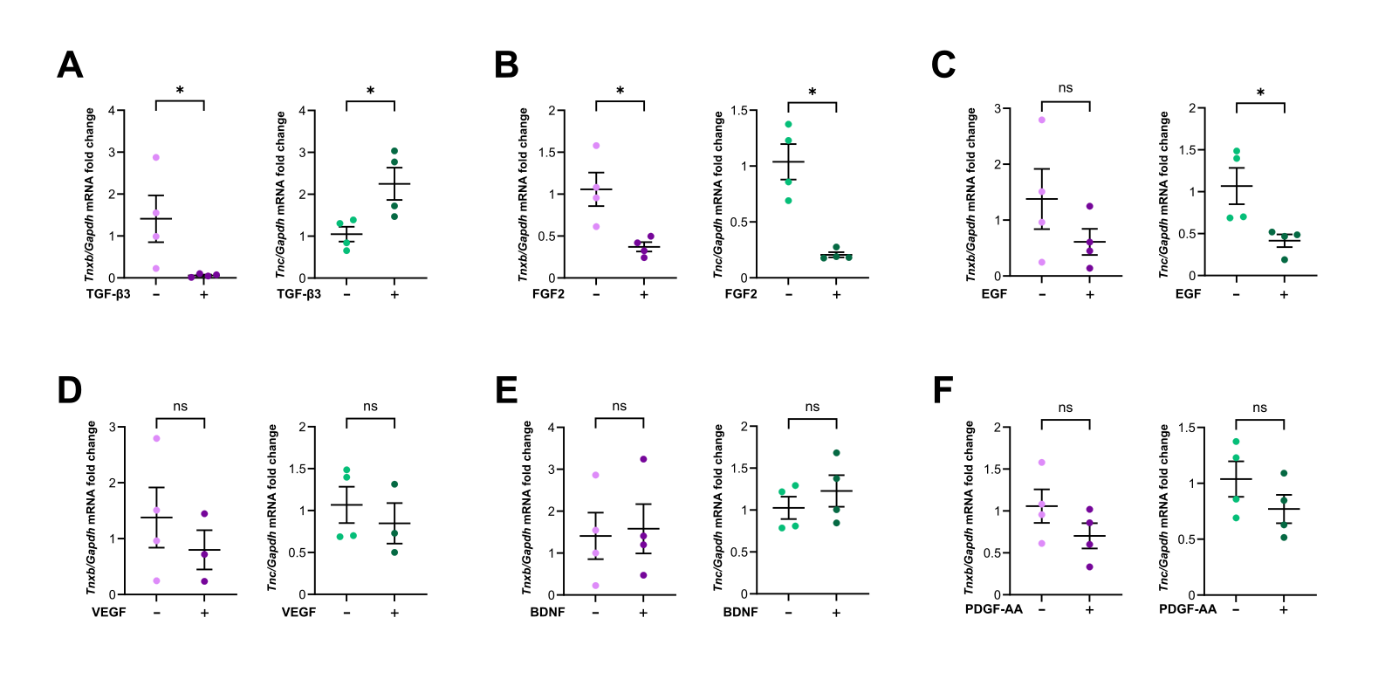
 ***Supplementary Figure S3: Effect of various selected growth factors on Tnxb and Tnc mRNA levels in KPC CAFs.*** *KPC CAFs were incubated without (-) or with (+) 20 ng/ml TGF-β3 (A),* FGF-2 (B), EGF (C), VEGF (D), BDNF (E) or PDGF-AA (F) *for 24h. Tnxb (left) and Tnc (right) mRNA levels were analyzed by qRT-PCR.*

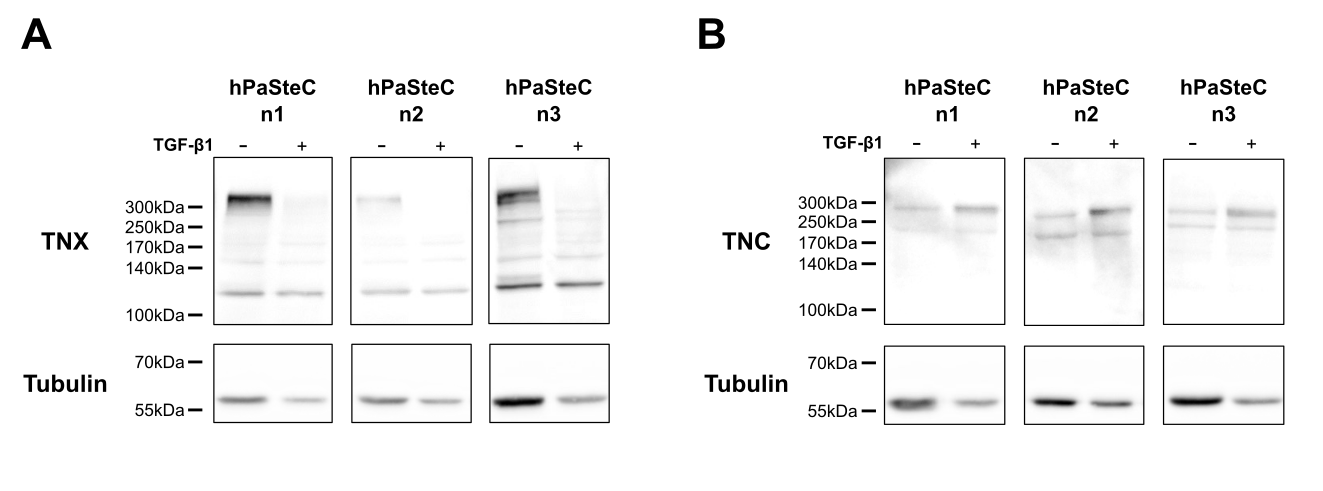

***Supplementary Figure S4: Regulation of TNX and TNC productions in human Pancreatic Stellate Cells (hPaSteC) following their stimulation with TGF-β1.*** *hPaSteC were serum-starved for 24h and then stimulated (+) or not (-) for 48h with 10 ng/ml TGF-β1. Protein extracts were analyzed by Western Blot for TNX (A), TNC (B) and acetylated Tubulin. 3 independent experiments (n) are shown.*

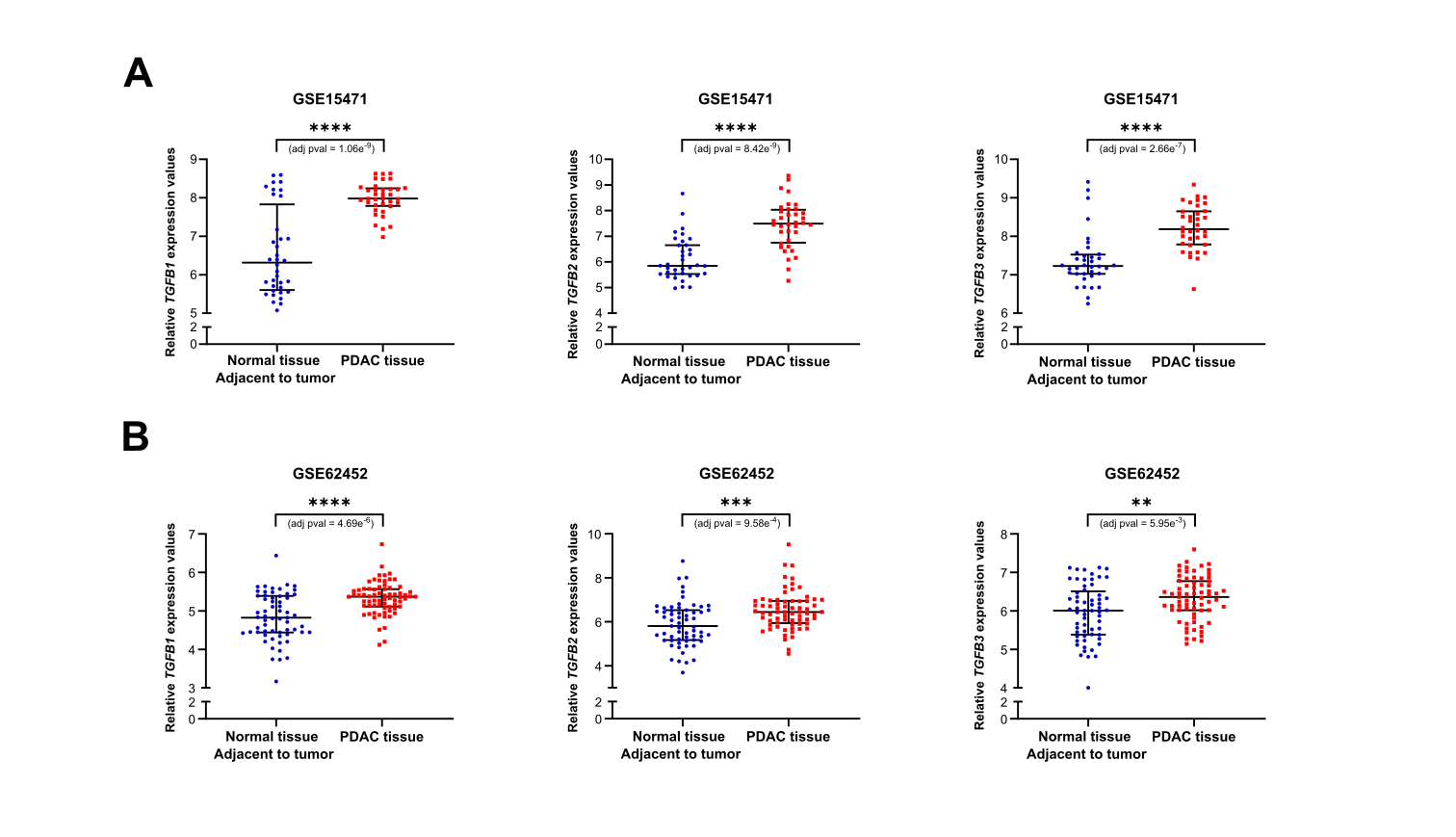
 ***Supplementary Figure S5: All members of the Transforming Growth Factor family are upregulated in PDAC.*** *Normalized* TGFB1 (left), TGFB2 (middle) and TGFB3 (right) *expression values in PDAC and NAT tissue samples for the GSE15471 (A) and GSE62452 (B) datasets (individual values with mean± SEM).*
